# Tissue Engineered Axon-Based “Living Scaffolds” Promote Survival of Spinal Cord Motor Neurons Following Peripheral Nerve Repair

**DOI:** 10.1101/847988

**Authors:** Joseph C. Maggiore, Justin C. Burrell, Kevin D. Browne, Kritika S. Katiyar, Franco A. Laimo, Zarina S. Ali, Hilton M. Kaplan, Joseph M. Rosen, D. Kacy Cullen

**Affiliations:** Department of Bioengineering, University of Pennsylvania, Philadelphia, PA, USA; Department of Neurosurgery, University of Pennsylvania, Philadelphia, PA, USA; Center for Neurotrauma, Neurodegeneration & Restoration, CMC VA Medical Center, Philadelphia, PA, USA; School of Biomedical Engineering, Science and Health Systems, Drexel University, Philadelphia, PA, USA; New Jersey Center for Biomaterials, Rutgers University, New Brunswick, NJ, United States of America, USA; Dartmouth-Hitchcock Medical Center, Division of Plastic Surgery, Dartmouth College, Lebanon, NH, USA

**Keywords:** Nerve regeneration, surgical repair, spinal cord motor neuron, neuronal survival, tissue engineered nerve graft

## Abstract

Peripheral nerve injury (PNI) impacts millions annually, often leaving debilitated patients with minimal repair options to improve functional recovery. Our group has previously developed tissue engineered nerve grafts (TENGs) featuring long, aligned axonal tracts from dorsal root ganglia (DRG) neurons that are fabricated in custom bioreactors using the process of axon “stretch-growth”. We have shown that TENGs effectively serve as “living scaffolds” to promote regeneration across segmental nerve defects by exploiting the newfound mechanism of axon-facilitated axon regeneration, or “AFAR”, by simultaneously providing haptic and neurotrophic support. To extend this work, the current study investigated the efficacy of living versus non-living regenerative scaffolds in preserving host sensory and motor neuronal health following nerve repair. Rats were assigned across five groups: naïve, or repair using autograft, nerve guidance tube (NGT) with collagen, NGT + non-aligned DRG populations in collagen, or TENGs. We found that TENG repairs yielded equivalent regenerative capacity as autograft repairs based on preserved health of host spinal cord motor neurons and acute axonal regeneration, whereas NGT repairs or DRG neurons within an NGT exhibited reduced motor neuron preservation and diminished regenerative capacity. These acute regenerative benefits ultimately resulted in enhanced levels of functional recovery in animals receiving TENGs, at levels matching those attained by autografts. Our findings indicate that TENGs may preserve host spinal cord motor neuron health and regenerative capacity without sacrificing an otherwise uninjured nerve (as in the case of the autograft), and therefore represent a promising alternative strategy for neurosurgical repair following PNI.

**HIGHLIGHTS:**

1. TENGs preserve host spinal cord motor neuron health and regenerative capacity acutely following repair of segmental nerve defects, matching that of the clinical gold-standard autograft and exceeding commercially-available nerve guidance tubes.
2. TENGs facilitated regeneration across segmental nerve defects, yielding similar degree of chronically surviving host spinal motor neurons and functional recovery as compared to autografts.
3. Early surgical intervention for segmental nerve defect with living scaffolds, such as TENGs and autografts, preserves the host regenerative capacity, and likely increases the ceiling for total regeneration and functional recovery at chronic time points compared to (acellular) commercially-available nerve guidance tubes.
4. TENGs preserve host neuronal health and regenerative capacity without sacrificing an otherwise uninjured nerve, and therefore represent a promising alternative strategy to autografts or nerve guidance tube repairs.

## Introduction

It is estimated that nearly 20 million Americans suffer from traumatic peripheral nerve injury (PNI), and out of those requiring surgical repair, only 50% achieve adequate functional recovery (Grinsell & Keating, 2014). Following severe PNI, the distal nerve segment undergoes Wallerian degeneration – a process of rapid axonal loss and myelin sheath degradation. During this process, contrary to degeneration, Schwann cells proliferate and form highly aligned columns called Bands of Büngner, which provide a pro-regenerative environment in the distal nerve that is necessary for axonal outgrowth from the injured proximal stump (Jessen & Mirsky, 2019). However, the pro-regenerative environment cannot be sustained in cases requiring long regenerative distances to the distal muscle and/or organ end-targets (Pfister et al., 2011). Indeed, prolonged denervation results in muscle atrophy and decreased receptiveness for reinnervation, diminished proximal neuronal health, retrograde dieback of axotomized neurons, and ultimately an overall lowering of regenerative capacity for functional recovery (Gordon, Tyreman, & Raji, 2011; Kaplan, Mishra, & Kohn, 2015; Pfister et al., 2011; Sakuma et al., 2016). As such, a PNI repair strategy that provides and sustains a pro-regenerative environment – including maintenance of proximal neuronal cell health – is necessary to establish a more comprehensive and effective clinical repair strategy.

In particular, one important cellular process involved with maintaining neuronal cell health is retrograde transport (Abe & Cavalli, 2008; Perlson, Maday, Fu, Moughamian, & Holzbaur, 2010). Following untreated PNI, retrograde transport of neurotrophic factors is inhibited in the proximal stump, concurrent with an initiation of a retrograde dieback process (Abe & Cavalli, 2008). Over time, chronically diminished retrograde transport leads to poor neuronal cell health, and reduced regenerative capacity of proximal neurons in the spinal cord (Terenghi, 1999). Thus, retrograde transport is a valuable surrogate marker for neuronal cell health and ultimately regenerative capacity. Recent studies have demonstrated that neurotrophic signals such as glial cell-line derived neurotrophic factor (GDNF), brain derived neurotrophic factor (BDNF) and nerve growth factor (NGF) secreted in the distal environment revitalize retrograde transport, improve cell health, and sustain regeneration (MacInnis & Campenot, 2002; Oudega & Hagg, 1996).

Recently, biological nerve grafts have been developed containing exogenous growth factors to simulate a pro-regenerative environment. However, a bolus of exogenous growth factors is unlikely able to provide a sustainable pro-regenerative environment required for functional recovery. Indeed, despite the recent advancements in biomaterials and tissue engineering, autograft nerve repairs still remain the current gold standard surgical treatment due to the superior functional recovery compared to commercially available products, such as a nerve guidance tube (NGT) or acellular nerve allograft. As an endogenously-available living scaffold, autografts provide regenerating host axons with anisotropic structural support, necessary for directed re-growth, as well as a rich supply of growth factors secreted by the Schwann cells (Kim, Yoo, & Atala, 2004). However, autografts are generally only partially effective for instances of large gap repairs and inherently cause donor site comorbidity.

In order to compensate for the drawbacks of current PNI repair strategies, our group has previously developed tissue engineered nerve grafts (TENGs) – generated utilizing stretch grown axons encapsulated in an extracellular matrix (ECM) that simultaneously provides structural support necessary for anisotropic growth and neurotrophic support through regulated growth factor release (Struzyna, Harris, Katiyar, Chen, & Cullen, 2015). These advantages likely play a significant role in the accelerated host nerve regeneration and enhanced functional recovery demonstrated in previous studies using TENGs (Huang et al., 2009; Katiyar et al., 2020; Pfister et al., 2011). While most PNI repair strategies have been well studied as they relate to regenerative and functional outcome measures (e.g., density of regenerating axons and extent of muscular recovery), there is limited information regarding the ability of various graft repair strategies to sustain proximal neuronal cell health and thereby maintain regenerative capacity. We hypothesize that living scaffolds, such as autografts and TENGs, will preserve proximal neuron health and overall capacity for regeneration by providing regenerating axons with haptic and chemotaxic cues, and/or trophic support (**Figure 1A**). Thus, we investigated the effect of various living (TENGs, dissociated neurons randomly distributed across ECM, and autografts) and non-living (ECM-laden nerve guidance tubes) scaffolds in maintaining the cell health of host spinal cord motor neurons and dorsal root ganglia neurons as a surrogate marker for regenerative capacity of motor and sensory axons, respectively. Within living scaffolds, this study also evaluated the effect of axonal organization, as highly organized TENGs with their architecture of defined neuronal populations spanned by long-projecting, aligned axonal tracts was compared to constructs comprised of less organized, dissociated neurons and axons growing randomly throughout the ECM.

**Figure 1.**
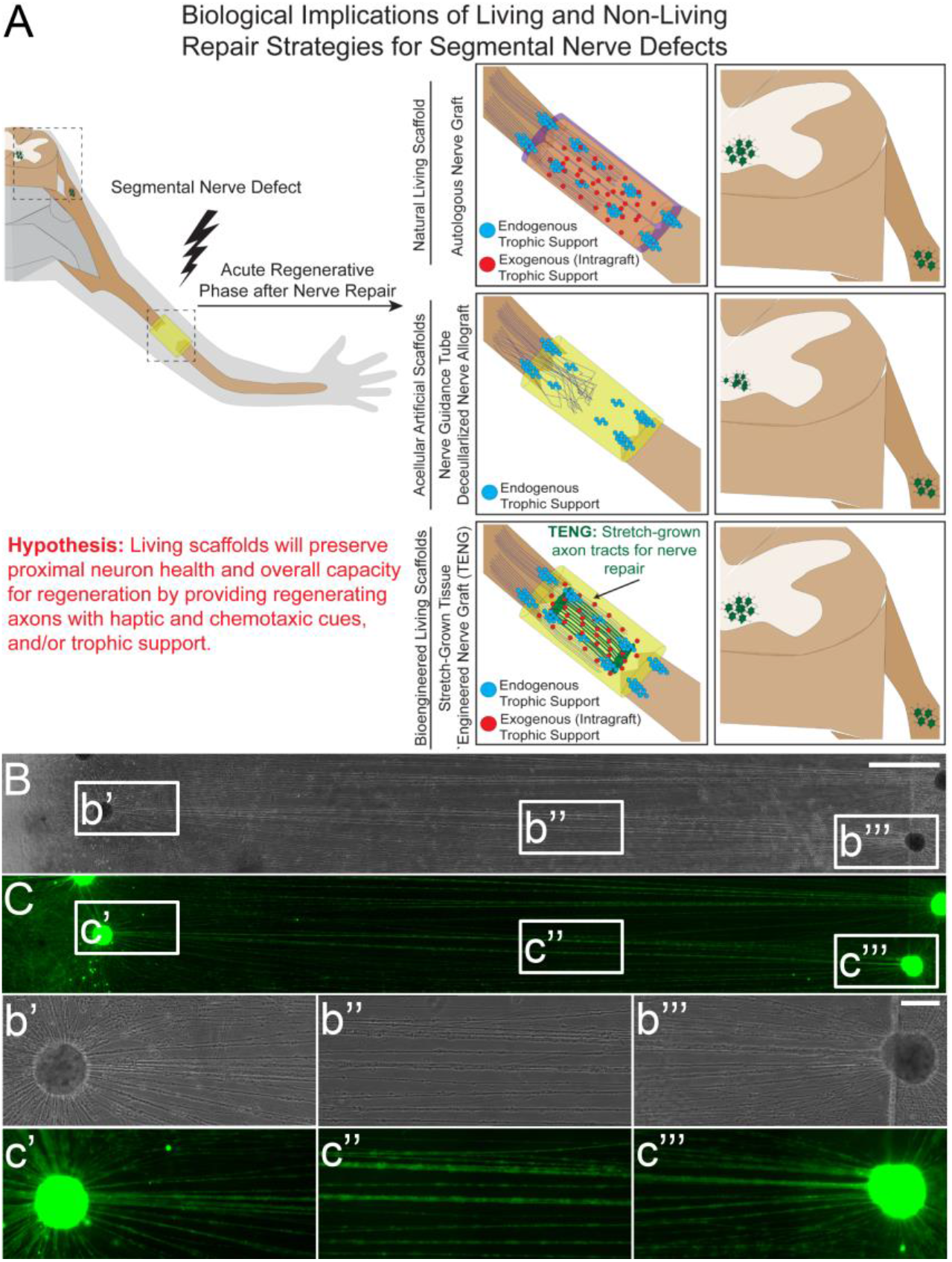
Biological Implications of Living and Non-Living Repair Strategies for Segmental Nerve Defects and Example of Axonal “Stretch-Growth” for Bioengineered Living Scaffold. (A) After segmental nerve injury, successful functional recovery necessitates that axons regenerate across a scaffold bridging the proximal and distal nerve stumps, enter the denervated distal nerve, and extend the entire distance to the end target. Often overlooked is that the surgical repair strategy may directly impact the ceiling for functional recovery. Autografts are natural living scaffolds that provide pro-regenerative trophic support, and chemotaxic and haptotaxic cues that enable accelerated axonal regeneration across the graft and is anticipated to preserve spinal motor neuron health, which would allow for greater functional recovery. Conversely, acellular strategies often lack neurotrophic and/or chemotaxic cues, leading to disorganized axonal growth, poor neuronal health, and ultimately diminished regenerative capacity. Our lab has developed tissue engineered nerve graft (TENGs) comprised of stretch grown axons that are bioengineered living scaffolds that preserve the ceiling for functional recovery by providing trophic support for regenerating axons, which we also postulate will preserve the regenerative capacity of the proximal neurons – a phenomenon that is the focal point of the current study. (B-C) To generate TENGs, embryonic rat GFP^+^ DRG explants were plated as two populations across a towing membrane separated by 500 μm. For 5 days *in vitro*, the DRG axons integrated and then were stretched to 1 cm with a micro-stepper over 6 days in a custom mechanobioreactor for transplantation. Example images are shown in (B) phase contrast and (C) and fluorescent microscopy. Once the axons reach the desired length, the TENG is encapsulated in collagen and rolled prior to transplantation. Scale bars: (Macro) 1000 μm; (Zoom in) 100 μm.

## Methods

All procedures were approved by the Michael J. Crescenz Veterans Affairs Medical Center Institutional Animal Care and Use Committee and adhered to the guidelines set forth in the NIH Public Health Service Policy on Humane Care and Use of Laboratory Animals (2015). All supplies were from Invitrogen (Carlsbad, CA), BD Biosciences (San Jose, CA), or Sigma-Aldrich (St. Louis, MO) unless otherwise noted.

### Dorsal Root Ganglion Neuron Isolation

Dorsal root ganglia (DRG) were isolated from embryonic day 16 Sprague-Dawley rats (Charles River, Wilmington, MA). Briefly, timed-pregnant rats were euthanized, and pups were extracted through Caesarian section. Each fetus was removed from the amniotic sac and put in cold Leibovitz-15. Embryonic DRG explants were isolated from spinal cords and either plated directly into media or dissociated. For dissociation, explants were suspended in pre-warmed trypsin (0.25%) + EDTA (1 mm) at 37°C for 45 min. Followed by the addition of neurobasal medium + 5% FBS, the tissue was vortexed for at least 30 seconds and then centrifuged at 1000 rpm for 3 minutes. The supernatant was aspirated, and the cells were resuspended at 5×10^6^ cells/mL in media. Media for both dissociated cells and explants consisted of neurobasal medium + 2% B-27 + 400-500 μm L-glutamine + 1% fetal bovine serum (Atlanta Biologicals) + 2.0-2.5 mg/mL glucose + 10-20 ng/mL 2.5S nerve growth factor, and a mitotic inhibitor formulation of 10 mM 5-fluoro-2’-deoxyuridine (5FdU) and 10 mm uridine to encourage non-neuronal cell elimination (Katiyar, Struzyna, Das, & Cullen, 2019).

### Nerve Repair Preparation

TENGs were fabricated by stretch-growing DRG explants plated in custom-fabricated mechanical elongation bioreactors as previously described (Huang et al., 2009; Katiyar et al., 2019; Katiyar et al., 2020; Pfister et al., 2006). Briefly, DRGs were plated in two populations along the interface of an aclar “towing” membrane treated with poly-D-lysine (20 ug/mL) and laminin (20 µg/ml), resulting in a separation of approximately 500 μm. Cells were transduced with an AAV viral vector (AAV2/1.hSynapsin.EGFP.WPRE.bGH, UPenn Vector Core) to produce GFP expression in the neurons. At 5 days *in vitro* (DIV) the cells were incubated overnight in media containing the vector (3.2×10^10^ genome copies/ml) and the cultures were rinsed with media the following day. Over 5 DIV, axonal connections were formed between the two populations. The populations were then gradually separated over the course of 6 days using a stepper motor system to displace the cells at a rate of 1 mm per day for 2 days and then 2 mm per day for 4 days, until the axons spanning them reached a total length of 1.0 cm as previously described (**Figure 1B-1C**). Fresh, pre-warmed media was used to replace the culture media every 2-3 days. After 11-13 DIV, once the stretched constructs had reached the desired length, their health was assessed. Constructs appropriate for transplant were removed from the incubation chambers and embedded in a collagen-based matrix (80% v/v) in Minimum Essential Media (MEM 10X) and sterile cell culture grade water supplemented with NGF (2.5S, 0.05 ng/mL). After gelation at 37°C, embedded cultures were gently removed and placed within a 1.2 cm long absorbable collagen nerve guidance tube (Stryker NeuroFlex™) and transplanted into a rat sciatic nerve injury model. For the NGT+DRG group, two populations of whole DRG explants (10 DRG in each row) were plated 1 cm apart on an ACLAR membrane to resemble the environment within the mechanobioreactor (Huang et al., 2009; Pfister et al., 2006). Axonal connections were allowed to form for 5 DIV as is done prior to initiation of mechanical tension for stretch grown constructs, and the cells were encapsulated in the collagen matrix described above and transferred into a 1.2 cm NGT for transplantation. The NGT control group received the same collagen matrix within the conduit.

### Surgical Procedure

Nerve regeneration was evaluated *in vivo* in a 1 cm rodent sciatic nerve injury model at 2 weeks following injury. A total of 20 male Sprague-Dawley rats were assigned to 5 groups: naïve (n=5), autograft (n=3), nerve guidance tube (NGT) (n=4), NGT containing disorganized DRG neurons (NGT+DRG; n=4), and TENGs (n=4). In addition, a total of 10 naïve rats were used to validate the fluorescent retrograde tracing methodology compared to conventional immunohistochemistry. Rats were anesthetized with isoflurane and the left hind was cleaned with betadine. Meloxicam (2.0 mg/kg) was given subcutaneously and bupivacaine (2.0 mg/kg) was administered along the incision line subcutaneously. The gluteal muscle was separated to expose the sciatic nerve exiting the sciatic notch. The sciatic nerve was carefully dissected to its trifurcation. The sciatic nerve was sharply transected 5 mm distal to the musculocutaneous nerve and a 1 cm nerve injury was made. For autograft repairs, a reverse-autograft technique was used (Roberts et al., 2017). For NGT, NGT+DRG, and TENG repairs, the nerve stumps were carefully inserted into each end of the nerve guidance tube with an overlap of 1 mm, and the epineurium was secured to the tube using 8-0 non-absorbable prolene sutures. The NGT+DRG repair contained DRG neurons embedded in collagen and NGT repairs contained collagen only as described above. The surgical site was closed with 3-0 absorbable vicryl sutures and skin staples. Animals were recovered and returned to the vivarium.

### Dye Application

At the terminal time point retrograde dye was applied to the nerve as previously described (Catapano et al., 2016). In brief, a nerve cuff was fashioned by capping silicon tubing (A-M Systems, 807600, 0.058” x 0.077” x 0.0095”) with PDMS (Fisher Scientific, Sylgard) that was trimmed to a length of 4 mm. The resulting nerve cuffs were stored in 70% ethanol until used. Prior to transplantation, the cuffs were rinsed in PBS and dried using a Kimwipe. In this study, retrograde dye transportation was evaluated following acute nerve regeneration at 14 days post repair (**Figure S1**). In a subset of animals, retrograde dye transportation was assessed following chronic nerve regeneration at 16 weeks post repair.

At 14 days after the initial repair procedure, the surgical site was re-exposed and the sciatic nerve was harvested, beginning 5 mm proximal to the repair zone, and immersion-fixed in formalin. A 2 mm by 2 mm piece of Kim wipe was soaked in 30% Fluoro-Ruby (FR; EMD Millipore, AG335) and placed inside the silicon nerve cuff toward the bottom. The silicon cuff was compressed, thereby creating a negative pressure vacuum, and the cuff was placed at the face of the proximal nerve stump. By slowly releasing the compression on the cuff, the negative pressure allowed for the sciatic nerve to seal within the cuff. Tisseel Fibrin Sealant (Baxter, 1504514) was applied around the cuff to fix the proximal nerve and cuff in place. The surgical site was closed using 3-0 absorbable vicryl sutures and skin staples, and the animal returned to the vivarium for 3 days of FR exposure (**Figure S1**).

### Nerve Regeneration and Schwann Cell Infiltration Across the Repair Zone

Formalin-fixed, frozen nerve sections were rinsed in PBS (3×5 min) and blocked for 1 hour in blocking solution (PBS with 4% normal horse serum and 0.3% Triton X-100). Primary antibodies diluted in blocking solution were then applied and incubated overnight at 4°C. Mouse anti-phosphorylated neurofilament (SMI-31, 1:1000, BioLegend Cat# 801601) and mouse anti-nonphosphorylated neurofilament (SMI-32, 1:1000, BioLegend Cat# 701601) were used to identify axons; rabbit anti-S100 Protein Ab-2 (S100, 1:500, Thermo Scientific Cat# RB-044-A) was used to identify Schwann cells. Following incubation, slides were rinsed in PBS (3×5 min) and secondary antibodies prepared in blocking solution were applied for 2 hours at room temperature: donkey anti-mouse 568 (Alexa Fluor® 568, 1:500, Thermo Scientific Cat# A10037) and donkey anti-rabbit 647 (Alexa Fluor® 647, 1:500, Thermo Scientific Cat# A31573). Sections were then rinsed in PBS (3×5 min), mounted with Fluoromount-G® (Southern Biotech Cat#0100-01) and coverslipped. Images were obtained with a Nikon A1R confocal microscope (1024×1024 pixels) with a 10x air objective and 60x oil objective using Nikon NIS-Elements AR 3.1.0 (Nikon Instruments, Tokyo, Japan).

### Spinal Cord Tissue Acquisition

Animals were transcardially perfused with 10% formalin and heparinized 0.9% NaCl. L4/L5 DRGs were extracted. Spinal cord T12-L6 region was extracted *en bloc*. All samples were fixed in paraformaldehyde overnight then placed in 30% sucrose for 48 hours. Full *en bloc* spinal cord was embedded in OCT then frozen. Tissue orientation was preserved with the use of tissue dye. Spinal cord samples were sectioned at 500 μm on a microtome cryostat and examined briefly under Nikon A1RSI Laser Scanning confocal microscope paired with NIS Elements AR 4.50.00 to screen sections with visible FR in the ventral horn. Spinal cord sections and DRGs with positive FR signal were stored in PBS for 24 hours.

### Spinal Cord Optical Clearing

Spinal cords were sectioned into 500 μm thick blocks and DRGs were optically cleared using the Visikol method and all washes were conducted at 15-minute time intervals unless otherwise stated (**Figure S2**). Spinal cord sections were washed in increasing concentrations of methanol (50%, 70%, and 100%) and stored for 12 hours in 100% methanol. Samples were then exposed to 20% DMSO/methanol followed by decreasing concentrations of methanol (80% and 50%). Next, samples were washed with PBS followed by PBS/0.2% Triton X-100. Next, samples were incubated in penetration buffer (0.2% Triton X-100, 20% DMSO, and 0.3M glycine in 1X PBS), then blocking buffer (0.2% Triton X-100, 6% NHS, and 10% DMSO in 1X PBS) at 37°C for 19 hours each. Samples were rinsed twice in washing buffer, then exposed to primary antibodies (1:500 Rabbit NeuN) in antibody buffer (0.2% Tween, 10 μg/mL Heparin, 3% NHS, and 5% DMSO in 1X PBS). After incubation in primary antibodies, samples were rinsed ten times in washing buffer (0.2% Tween and 10 μg/mL Heparin in 1X PBS), followed by exposure to secondary antibodies (1:500 Donkey anti-rabbit 647). Samples were again rinsed ten times in washing buffer. Finally, samples were exposed to increasing concentrations of methanol (50%, 70%, and 100%) at three rinses each while incubated at 37 °C. The samples were then incubated in Visikol 1 for 24 hours, then Visikol 2 for 48 hours. For all DRG samples, these samples were placed in Visikol 1 after the initial ascending methanol steps.

### Quantification of Ventral Horn and Dorsal Root Ganglia

For conventional immunohistochemistry: naïve spinal cords were harvested 3 days following FR application in order to validate the optical clearing technique. Briefly, samples were stored in 30% sucrose for 24 hours or until saturation for cryoprotection, embedded in OCT, and frozen in −80 **°**C isopentane. Axial sections were taken using a cryostat microtome (30 μm thick) and stained for NeuN (1:500 Rabbit NeuN, 1:500 donkey anti-rabbit 647). Slides were imaged at 20x using a Nikon A1RSI Laser Scanning confocal microscope paired with NIS Elements AR 4.50.00, taking z-stacks at 5 μm intervals. FR and NeuN cell counts were quantified from maximum projections. The Abercrombie correction for cell quantification was applied based on section thickness and an estimated cell size of 30 μm (Abercrombie, 1946).

For optical cleared tissue: samples were placed in glass bottom well plates and immersed fully in Visikol 2. Tissue was then imaged using Nikon A1RSI Laser Scanning confocal microscope under 10x air objective. All images were taken using a z-stack at 5 μm steps with laser settings optimized for naïve tissue. Images were analyzed using ImageJ FIJI software. Maximum projections of 100 μm were generated and all cells within the ventral horn (L4-L6) of the spinal cord and DRG (L4, L5) were quantified by drawing line ROI through background and the entirety of the cell body, which was inspired by previous methodology (Cullen, Vernekar, & LaPlaca, 2011). Using a custom MATLAB script, the intensity value for each cell body was calculated with the following formula:

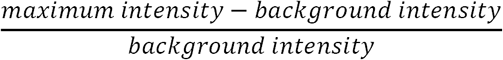

### Functional Assessment

At 16 weeks post repair, compound muscle action potential (CMAP) was assessed to evaluate functional regeneration. Animals were re-anesthetized and the graft was re-exposed, A bipolar subdermal electrode was placed in the tibialis anterior and the nerve was stimulated (biphasic; amplitude: 0–10 mA; duration: 0.2 ms; frequency: 1 Hz) using a handheld bipolar hook electrode (Rochester Electro-Medical, Lutz, FL; #400900) 5 mm proximal to the repair zone. The supramaximal CMAP recording was obtained and averaged over a train of 5 pulses (100x gain and recorded with 10–10,000 Hz band pass and 60 Hz notch filters).

### Statistical Analysis

For conventional and optical cleared quantification: the mean and standard error were calculated for cell counts and compared using a parametric one-way ANOVA with multiple comparisons. To compare the number of FR^+^ cells across each experimental group, a parametric ANOVA test with multiple comparisons was conducted with an alpha value of 0.05. The intensity of every cell per animal was log transformed to fit the naïve group data to a normal distribution. Frequency distributions for each experimental group were calculated with a bin length of 0.1 for MN and DRG. These frequency distributions were normalized by dividing each bin frequency by the experimental group sample size. The mean intensity of each animal was averaged, and experimental groups were compared using a nonparametric Kruskal-Wallis ANOVA with multiple comparisons. A linear regression was generated between number of FR^+^ cells and the transformed mean intensity value.

## Results

### Repair Zone Analysis

A 1-cm segmental defect was created in the sciatic nerve of rats, with immediate surgical repair using a reverse-autograft, an NGT containing a collagen-based ECM, an NGT containing disorganized DRG neurons throughout the ECM (“NGT+DRG”), or custom-fabricated TENGs featuring longitudinally-aligned axonal tracts projecting from DRG neurons within ECM. To evaluate acute regeneration processes across the repair zones, at 2-weeks post-repair longitudinal sections from rat sciatic nerve grafts were immunostained with SMI31/32 and S-100 to visualize axons and Schwann cells, respectively, similar to previous studies (Katiyar et al., 2020). Axonal infiltration from the proximal region into the graft region was observed across all groups (**Figure 2A-2D**). Regenerating axons and Schwann cell infiltration were observed at higher magnification in longitudinal sections of the graft region. Implanted DRG expressing GFP survived transplantation in the NGT+DRG group (**Figure 2C**). Moreover, TENG neurons also survived transplantation and extended axons into the distal nerve, and integrated with host Schwann cells and axons, enabling axon-facilitated axonal regeneration (AFAR) across the graft zone (**Figure 2D**).

**Figure 2.**
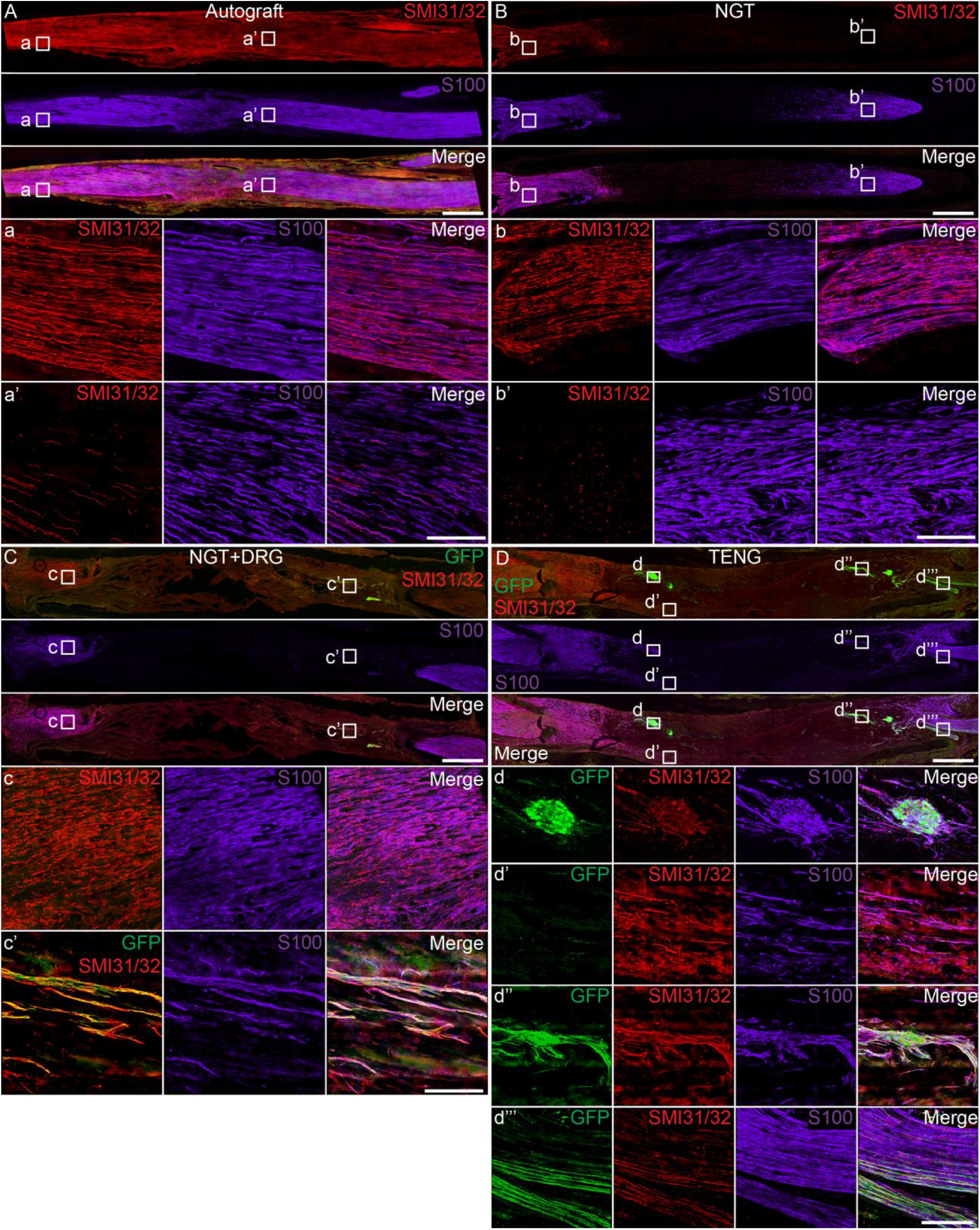
Host Axon Regeneration and Schwann Cell Infiltration at 2 Weeks Following 1 cm Lesion in Rat Sciatic Nerve. Confocal reconstruction of longitudinal frozen sections (20 μm) of a 1 cm rodent sciatic nerve segmental repair using (A) autograft, (B) NGT, (C) NGT+DRG, and (D) TENG. (a-d) Higher magnification images showing sections labeled for regenerating axons (SMI31/32^+^) and Schwann cells (S100). Transplanted DRG neurons expressing GFP were observed in the repair zone of the (C) NGT-DRG and (D) TENG groups. (A, D) Robust host axonal ingrowth and Schwann cell infiltration was observed across the autograft and TENG. Comparatively, axon extension and Schwann cell infiltration were attenuated following repair with (B) NGT. (C, D) Transplanted DRGs survived and facilitated host axon across the graft. (d, d’’) Higher magnification images revealed TENG neuronal survival post transplantation and close integration with host Schwann cells and axons. (d’) Host axons extending across the repair zone were often found near TENG axons spanning the length of graft. (d’’’) TENG and host axons were also observed extending into the distal nerve. Scale: A-C 1000 μm; a-c 100 μm.

### Neuronal Labeling Validation in Optically Cleared Sections from Naïve Tissue

To validate the optical clearing technique and quantification methodology used in subsequent analyses, FR and NeuN were visualized in 500 μm thick naïve spinal cord blocks. FR labeled cells were clearly visualized within optically cleared spinal cord stained for NeuN in both axial and longitudinal blocks and adequate antibody penetration was confirmed in the volumetric reconstruction of the Z-stack images across various X-Y-Z planes **(Figure 3**). FR and NeuN cells were quantified in maximum projection z-stack images for 500 μm thick sections of spinal cords and 30 μm thick frozen sections. Conventional IHC cell counts were significantly higher for FR and NeuN counts than the IHC quantification with the Abercrombie correction (IHC+AC) or Visikol HISTO sectioning. No statistical significance was found for FR or NeuN quantification between the IHC+AC and Visikol HISTO methods (p > 0.05).

**Figure 3.**
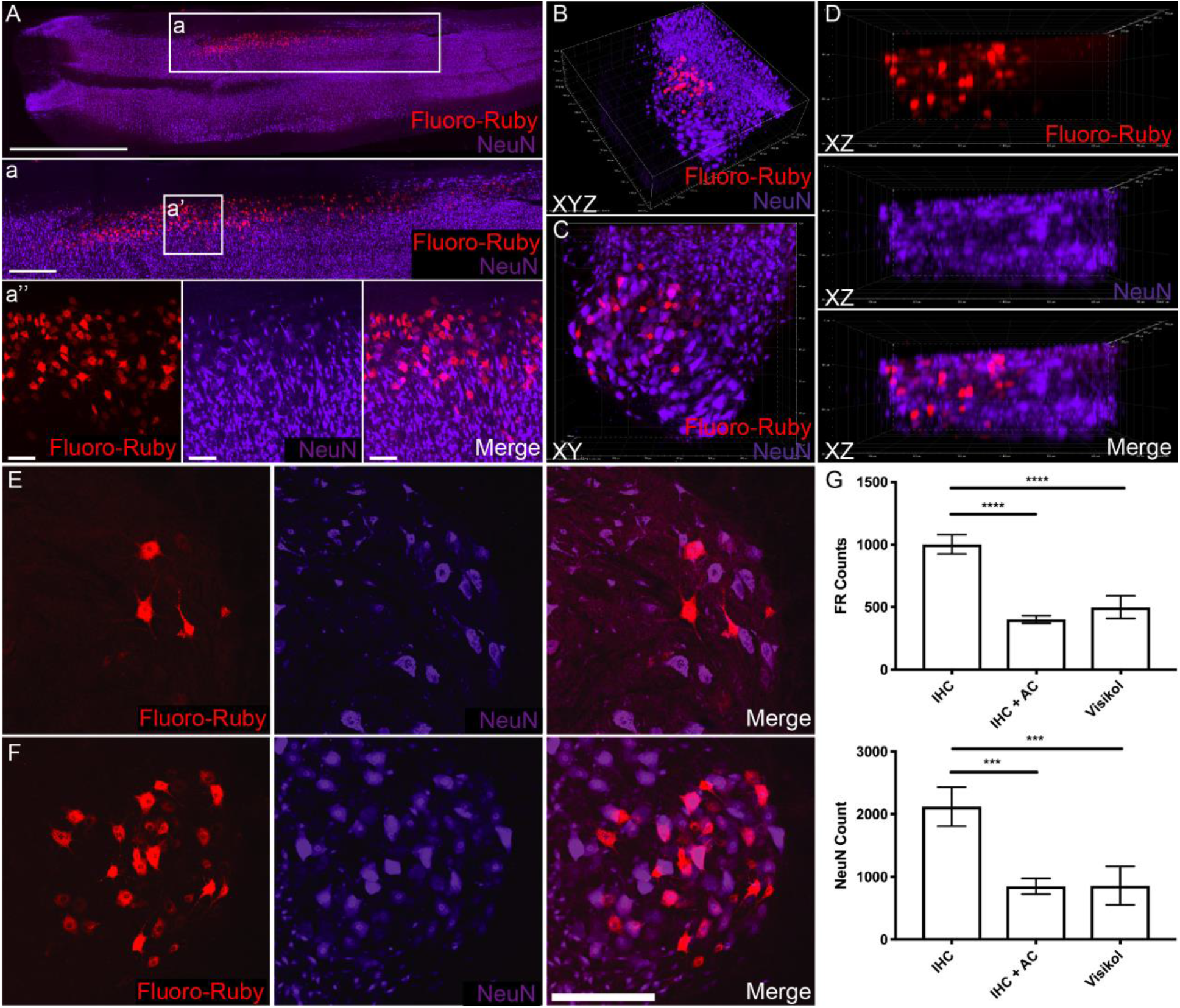
FR and NeuN Quantification in Naïve Spinal Cord Tissue to Validate Optical Clearing Technique with Conventional IHC Methodology. Visikol HISTO protocol can be used to analyze spinal cord tissue longitudinally or axially in 500 μm thick macro sections. (A) Fluoro-Ruby labelled cells and NeuN immunostained cells can be visualized in maximum projection z-stacks. Scale bars: 3000 μm, 600 μm, 60 μm. (B) NeuN antibody penetration was visually confirmed through the complete 3D confocal reconstruction. XYZ view of axial spinal cord ventral horn section labelled with Fluoro-Ruby and NeuN. (C) XY 3D view of axial spinal cord ventral horn section. (D) XZ view of axial spinal cord section, confirming sufficient laser penetration to view Fluoro-Ruby through entirety of section. (E) Standard IHC (30 μm histological samples), z-stack maximum projection. (F) Spinal cord tissue blocks (500 μm sections) stained for NeuN IHC during Visikol HISTO process, z-stack maximum projection. Scale bar: 300 μm. (G) No statistical significance was found between 30 μm sections stained using conventional IHC with Abercrombie correction (IHC+AC) and Visikol cleared tissue. Error bars represent standard deviation. *** = p<0.001; **** = p<0.0001.

### Ventral Horn Retrograde Labeling Analysis Following PNI Repair

The retrograde dye FR was applied to the nerve segment proximal to the repair zone at 14 days post-repair, and the presence of FR in proximal host neurons was assessed 3 days thereafter as a surrogate for neuronal health and active regeneration. In the spinal cord, TENG repairs exhibited a similar number of FR^+^ labelled MN cell bodies (1038.0 ± 100.5) as naïve animals (935.0 ± 35.4) and autograft repairs (914.7 ± 35.4) (p > 0.05) (**Figure 4**). TENG repairs also exhibited a significant increase in FR^+^ MNs compared to NGT (357.3 ± 52.3, p < 0.001) and NGT+DRG repairs (678.8 ± 82.6, p < 0.05). NGT repairs exhibited a significant decrease in the number of FR^+^ MN cell bodies as compared to naïve, autograft and TENG repairs (p < 0.001 each), as well as NGT+DRG repairs (p < 0.05) (**Figure 4F**). The density of NeuN^+^ was also quantified within the ventral horn of the spinal cord. NGT NeuN counts were significantly less than naïve and autograft NeuN counts (p < 0.05), while NGT+DRG and TENG NeuN counts were not statistically different from naïve animals (p > 0.05) (**Figure 4G**). The intensity of FR uptake per neuron was also quantified as a further metric of neuronal health. NGT repairs exhibited a significantly diminished number of high FR intensity cells than the naïve group. Autograft repairs showed a rightward shift in FR intensity as compared with naïve group while TENG repairs showed a similar FR intensity profile as naïve repairs (**Figure 4H;** See **Figure S3** for a breakout of various distributions per experimental group).

**Figure 4.**
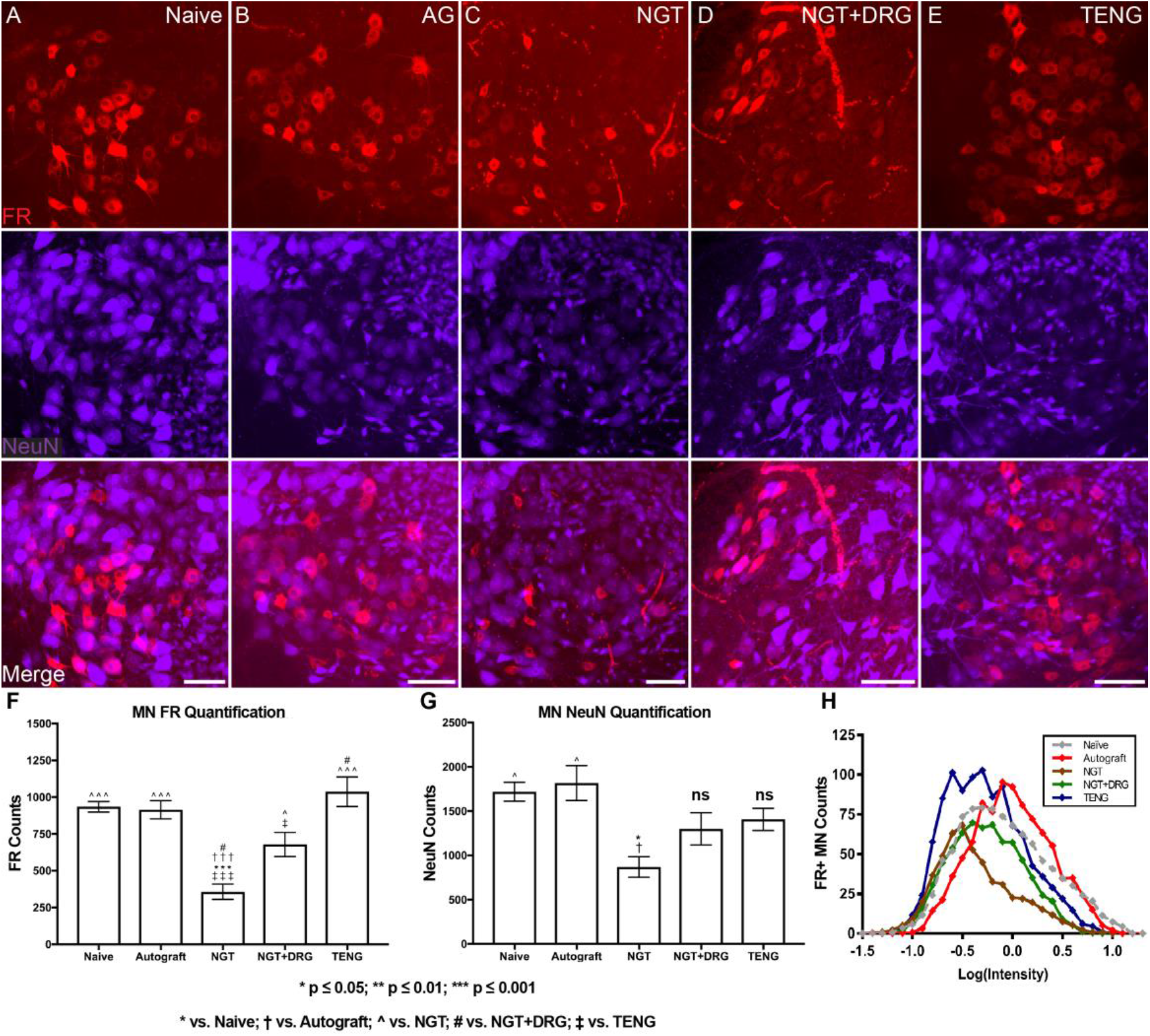
FR Retrograde Tracing and MN Quantification in the Ventral Horn Spinal Cord. (A-E) MN cell bodies labeled with FR and NeuN were visualized in the ventral horn of optically cleared spinal cords. (F) FR^+^ MN cell bodies in the ventral horn were quantified. When compared with naïve, both NGT (p < 0.0001) and NGT+DRG (p < 0.05) repair strategies produced a statistically significant decrease in mean count. (G) NeuN cell bodies were quantified in the ventral horn and a statistically significant reduction was observed following the NGT repair group compared to naïve (p < 0.05). Scale bar: (Macro) 150 μm; (Zoom in) 150 μm. Error bars represent SEM. ns denotes no significance compared to naïve. (H) Frequency distributions of MN FR fluorescence intensity were plotted. Fluorescence intensity was calculated for each FR^+^ cell in the ventral horn by measuring the maximum intensity of the cell relative to the local background. Individual cell intensity was log transformed to fit a normal distribution. See **Figure S3** for frequency distribution profiles for each experimental group relative to the naïve frequency distribution.

### Dorsal Root Ganglia Retrograde Labeling Analysis Following PNI Repair

Similar to the MN analysis of retrograde FR labeling, the density and intensity of FR labeling in L4 and L5 DRG were also assessed. There were no statistical differences in the number of FR^+^ cells across experimental groups in the L4 and L5 DRG regions (**Figure 5**). The intensity of FR uptake per neuron was again quantified as a metric of neuronal health. Autograft repairs exhibited a similar FR intensity profile as naïve animals, while NGT, NGT+DRG and TENG repairs showed a downward shift in FR intensity profile as compared with the naïve group (**Figure 5H;** See **Figure S4** for a breakout of various distributions per experimental group).

**Figure 5.**
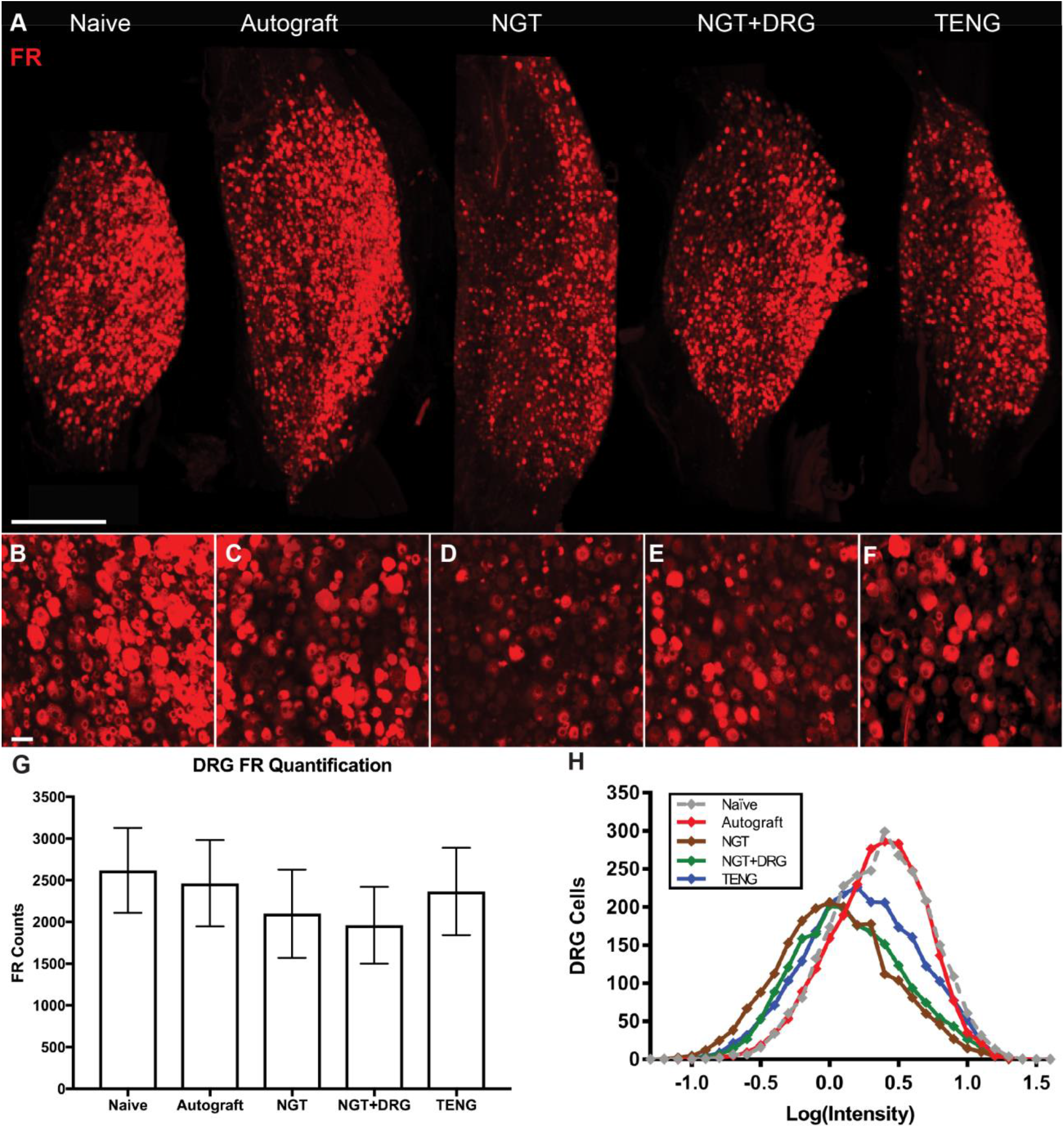
FR Quantification in L4 and L5 DRG. L4/L5 DRG samples were visualized *en bloc* following whole mount optical tissue clearance. (A-F) Representative confocal reconstructions generated from z-stack max projections are shown. (G) No statistical difference was observed between any treatment group for L4 and L5 DRG FR counts. Error bars represent SEM. Scale bar: (A) 700 μm; (B-F) 50 μm. (H) Frequency distributions of DRG fluorescence intensity were plotted. Fluorescence intensity was calculated for each FR^+^ cell in the ventral horn by measuring the maximum intensity of the cell relative to the local background. Individual cell intensity was log transformed to fit a normal distribution. See **Figure S4** for frequency distribution profiles for each experimental group relative to the naïve frequency distribution.

### FR Uptake and Functional Assessment at 16 Weeks Following PNI Repair

Structural and functional measures were assessed at 16 weeks following PNI repair to relate acute changes in retrograde neuronal labeling with more chronic outcomes. A qualitative assessment of a subset of animals at 16 weeks post repair revealed similar numbers of FR labeled spinal motor neurons following TENG and autograft repair, as compared to a reduced number of labeled neurons following NGT repair (**Figure 6**). No differences were observed in the labeling of DRG neurons between groups. CMAP recordings were obtained in all animals at 16 weeks post repair, revealing that the amplitude of the response was consistently much greater following TENG or autograft repair than following NGT repair. This observation matches our previously reported quantitative assessment of functional recovery using these repair groups (Katiyar et al., 2020). The inclusion of these functional recovery measures in the current study provides further context and a neurobiological mechanism that likely at least partially explains decreased CMAP following an acellular repair strategy as compared to enhanced CMAP following living scaffold repair strategies (i.e. autografts and TENG repairs).

**Figure 6.**
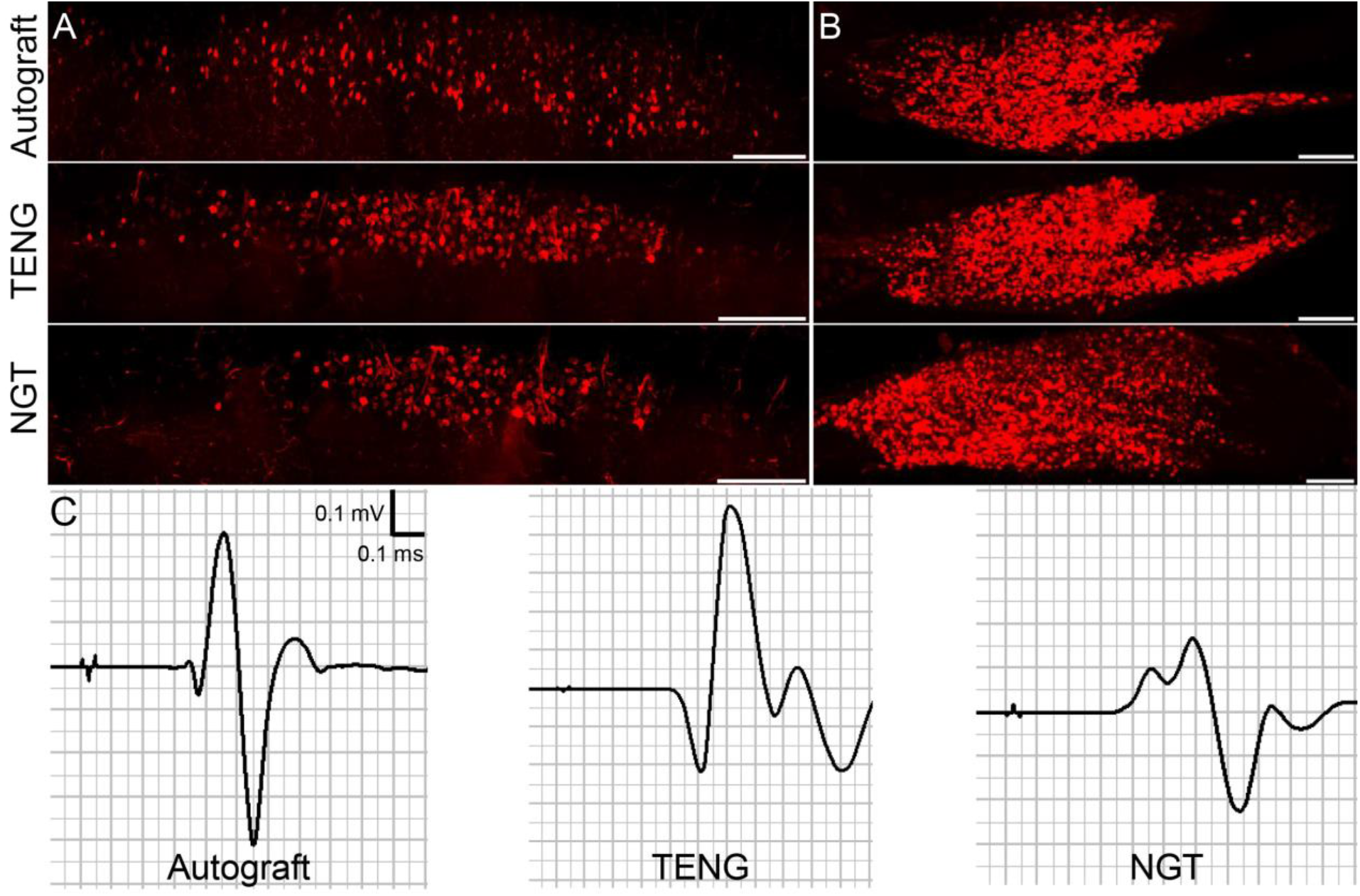
FR Uptake in Spinal Motor Neurons and DRGs at 16 Weeks Post Repair. In a subset of animals, (A) spinal motor neurons and (B) DRGs were labeled with FR at 16 weeks following peripheral nerve repair. Qualitative assessment revealed similar number of labeled spinal motor neurons following TENG and autograft repair and no differences were observed in the DRG between groups. (C) Similar muscle electrophysiological recovery was observed between the TENG and AG repair groups. Scale bar: 100 μm.

## Discussion

PNI recovery is a race against time since regenerating axons have a limited period to reach distal end targets before the pro-regenerative distal nerve environment loses its capacity to support regeneration. Here, various approaches have been developed to facilitate nerve regeneration across a graft region and ultimately sustain the pro-regenerative environment in the distal segment (Patel, Lyon, & Huang, 2018). However, there has been limited focus on the ability for a graft to influence the intrinsic regenerative capacity of proximal neurons. To address this gap, the current study compliments our previous efforts for TENG development and implementation by highlighting an additional mechanism by which this class of tissue engineered “living scaffolds” may lead to an improved time course and extent of functional recovery. Of note, this mechanism of preserving proximal neuronal health and regenerative capacity was superior using TENGs as compared to acellular repair strategies, such as commonly used NGTs, and resulted in a commensurate level of increased functional recovery. This finding builds on a previous recent report, where we showed that TENGs enable “axon-facilitated axonal regeneration or AFAR” resulting in accelerated axonal regeneration and functional recovery compared to commercially-available NGTs alone in a similar rat model of PNI (Katiyar et al., 2020). To the best of our knowledge, this is the first study to investigate the effect of different peripheral nerve repair strategies on proximal neuron health and regenerative capacity.

In this study, FR expression in host spinal cord motor neurons and DRG neurons was quantified as a surrogate marker for retrograde transport capability and overall neuron health. At 2 weeks post repair, a similar number of FR^+^ cells was found in the ventral horn in animals repaired using TENGs or autografts, and both matched that found in naïve animals. In contrast, NGT repairs resulted in >65% reduction in FR^+^ neurons as compared to TENG repairs, likely due to acellular NGTs not providing adequate neurotrophic and anisotropic structural support. Furthermore, the average fluorescent intensity of the FR^+^ spinal MNs in the NGT repair group was significantly less than that found in naïve animals. In addition, nerves repaired with NGTs seeded with disorganized DRG populations exhibited a modest increase in the level of retrograde transport compared to the NGT group, potentially indicating the benefit of a cellular-derived trophic component alone for PNI repair. The benefit of a living cellular component in a nerve graft was further corroborated with evidence of Schwann cell infiltration surrounding transplanted cells within the graft zone. Although the reduction in FR^+^ labeling in the ventral spinal cord of the NGT group compared to NGT+DRG groups indicates the advantage of living cells within the graft, the extent of FR^+^ cells in the NGT+DRG group was significantly less that that found following TENG or autograft repairs. This suggests the dual importance for nerve grafts to supply trophic factors while simultaneously presenting a biomimetic, anisotropic structural organization.

This study also assessed the presence of NeuN^+^ neurons in the spinal cord and DRG following various PNI repair strategies. Historically, NeuN has been considered a reliable mature neuron marker that is highly conserved across species and useful for quantifying neuronal density (e.g., to indirectly assess neuronal loss). However, recent work has shown that NeuN antibody specificity generally depends on phosphorylation state. Anti-NeuN binds to phosphorylated NeuN within neuronal nuclei, therefore, loss of NeuN immunoreactivity may indicate a stress response but not necessarily neuronal death (Duan et al., 2016). In this study, NeuN counts in the spinal cord were similar between naïve animals and autograft repairs, and neither were significantly different from TENG or NGT+DRG repairs. In contrast, fewer NeuN-labeled spinal MNs were visualized following NGT repair. The implication of these findings is unclear as NeuN expression has not been well characterized with respect to injury severity or repair strategy. However, facial nerve axotomy has been reported to result in rapid and protracted decrease in NeuN expression compared to transient NeuN loss after facial nerve crush injury (McPhail, McBride, McGraw, Steeves, & Tetzlaff, 2004). In addition, chronic nerve axotomy induced by avulsion injury was shown to result in a rapid and persistent loss of NeuN expression in the spinal cord ventral horn; however, motor neuron death progressed over the course of 4 weeks (Tan et al., 2015). The rapid loss of NeuN expression at this acute time point is likely associated with biochemical disturbances and molecular changes to the injured cell body resulting from the axonal disconnection to the distal structures and deprivation of Schwann cell- and/or muscle-derived neurotrophic support (Tan et al., 2015). Therefore, injury severity and grafting strategy may have a major role in NeuN labeling an/or expression.

Whether the loss of NeuN expression is persistent or transient may be dependent on the injury severity. After crush nerve injury, the spared underlying nerve tissue provides a robust cache of neurotrophic support from the distal structures, enabling more rapid regeneration to the end target. In contrast, severe nerve injury, such as chronic nerve axotomy or root avulsion, deprives the cell body of crucial neurotropic factors secreted by the distal structures (Fu & Gordon, 1995; Funakoshi et al., 1993; Gordon, 2009; McPhail et al., 2004; Peyronnard, Charron, Lavoie, & Messier, 1986; Tan et al., 2015; Wiberg, Kingham, & Novikova, 2017). These mechanisms may also play a significant role in differences in NeuN expression after segmental nerve repair. Following autograft repair, trophic factors secreted by Schwann cells within the donor nerve may help to sustain the injured proximal neurons. In contrast, NGTs lack endogenous Schwann cells, which may deprive the proximal neurons of neurotrophic factors and result in decreased NeuN expression. Following NGT repair, acute loss of NeuN expression could be transient and reverse as the regenerating axons interact with pro-regenerative Schwann cells moving in from the distal nerve stump; however, our findings at 16-weeks post repair suggest that this NeuN loss may be permanent, and thus represent neuronal cell death. In contrast, similar to autografts, the DRG+NGT and TENG groups likely preserved NeuN expression by providing regenerating axons with trophic factor support; however, the exact mechanism remains unclear as these constructs were not supplemented with Schwann cells. Overall, our findings warrant follow-on studies to determine whether decreased NeuN expression following segmental nerve defects provides an indication of neuronal cell health or if loss of expression is generally associated with permanent neuronal cell death. If the decreased NeuN expression is transient, then these findings potentially suggest strategies lacking trophic support may result in a large number of unhealthy proximal neurons during a crucial period for regeneration. Moreover, the ability for living scaffolds to sustain neuronal health at an acute time point further suggests the importance for maintaining overall regenerative capacity to enhance the rate and extent of functional recovery.

Although there were stark differences in the level of FR and NeuN expression in the ventral spinal cord, there did not appear to be any significant change in DRG neurons based on FR or NeuN expression across any of the experimental groups. These results are consistent with previous studies that found no change in the number of retrogradely labeled DRG neurons at two weeks following sural nerve axotomy (Peyronnard et al., 1986). Previous studies also corroborate this finding, demonstrating that sensory neurons are more resilient and regenerate more robustly than motor neurons at early time points following nerve axotomy (Cheah, Fawcett, & Haenzi, 2017). More extensive studies are necessary to elucidate the intrinsic gene expression patterns and physiological pathways in DRG that make these neurons less dependent in extrinsic factors for injury and regeneration responses.

In the current study, retrograde transport was used as a surrogate marker for neuronal cell health and potential regenerative capacity, with quantitative measurements made of the density of FR^+^ cells and well as the per-cell intensity of FR expression. Indeed, retrograde signaling is necessary to increase protein synthesis and enable growth cone extension (Rishal & Fainzilber, 2010). While there is substantial evidence that retrograde transport is modulated through varying levels of neuronal cell health (Perlson et al., 2010), other mechanisms, such as the rate of axonal resealing, might modulate the amount of FR transported to the neuronal cell body (Howard, David, & Barrett, 1999). Previous studies have shown evidence that axon diameter may also influence rate of microtubule-based retrograde transport (Xia et al., 2003). In order to conclusively implicate active retrograde transport as the modulator of observed changes in FR expression, it may be of interest in future experimental designs to include a control repair group that is exposed to a retrograde transport inhibitor, such as Ciliobrevin D (Firestone et al., 2012). In addition, the per-cell intensity of FR expression was used as a representative marker of the extent of retrograde transport and therefore as an indirect marker of neuronal health. Of note, the fluorescent intensity of the cell body was calculated using the subtraction of the background intensity from the maximum intensity relative to the background intensity based on previous methodology (Cullen et al., 2011). However, in future analyses, alternative methodology such as comparing the mean pixel intensity across cell bodies and/or factoring in the somata volume may offer additional useful information on neuronal health.

Living scaffolds may provide a sustained bolus of numerous pro-regenerative neurotrophic factors that support regenerating axons (and hence the neurons that project these axons) and ultimately facilitate functional recovery. In this study, we found TENGs and autografts had a similar degree of healthy neurons at an acute time point post-injury and similar levels of functional recovery chronically. Both autograft and TENG repairs resulted in substantially improved metrics of acute regeneration and chronic recovery than NGT repairs. It is important to note that these results should be interpreted in context, as some metrics were not statistically significant yet there were large differences in means, suggesting that although these findings are promising, further optimization may improve the consistency of improvements in neuronal health and ultimately functional recovery. In addition, future studies are necessary to more directly investigate whether early recovery of neuronal health also improves the capacity for muscle reinnervation. To further understand regenerative capacity for clinical applicability, future studies might also include additional cell markers to provide additional insight into the neurons that are actively regenerating toward the end targets. The varying expression profile of certain transcriptional factors, such as ATF-3 and C-JUN, following nerve injury, during regeneration, and until reinnervation could be combined with FR expression data to provide greater insight into the duality between regeneration and neuronal cell health (Stenberg, Kanje, Dolezal, & Dahlin, 2012).

Although TENGs have demonstrated the potential for preserving neuronal cell health, further optimization is necessary to tailor the repair strategy for specific injuries. For example, sensory nerve autografts are typically used to repair all injuries, including primarily motor as well as mixed motor-sensory nerves. However, there has been some evidence that suggests motor nerve autografts may further increase functional recovery (Nichols et al., 2004). Therefore, it might be useful to develop modality-specific TENGs comprised of sensory, motor, or mixed motor-sensory neurons/axons to further enhance the regenerative capacity across multiple types of nerves. However, in this study, TENGs comprised of DRG neurons/axon tracts were shown to maintain neuronal cell health in MN regions of the spinal cord.

To date, most commercially available NGTs are empty conduits which lack neurotrophic and anisotropic support (e.g., NeuroTube^®^, poly-glycolic acid, Baxter/Synovis; NeuraGen^®^, collagen, Integra LifeSciences; Neuroflex™, collagen, Stryker), although there has recently been the introduction of an NGT filled with collagen containing aligned channels to provide anisotropic guidance (i.e., Nerbridge^®^, Toyobo). Numerous previous studies have reported that repairs using an NGT resulted in diminished functional recovery compared to an autograft repair, even in gap repairs less than 1.5 cm (Kaplan et al., 2015). Our study corroborated this by demonstrating that an acellular graft with solely structural isotropic support from collagen was unable to support proximal cell survival and regenerative capacity as compared to other grafts. NGTs are therefore inadequate clinical vessels for supporting effective, healthy, and functional regeneration in all cases. Acellularized nerve allografts (ANAs) were developed as an alternative non-living scaffold repair strategy that provided regenerating axons with the nerve architecture and haptic cues similar to an autograft repair (Moore et al., 2011). However, ANAs lack a cellular component and the necessary trophic support for sustained nerve regeneration, thus likely minimizing regenerative capacity and the ultimate level of functional recovery (Tang et al., 2013).

Recent efforts have aimed to develop biomimetic materials and controlled release systems to offer alternative strategies to acellular NGTs by adding biologically-active structural proteins and soluble factors, such as ECM, NGF, BDNF, and GDNF (Ramburrun et al., 2014; Wieringa, Goncalves de Pinho, Micera, van Wezel, & Moroni, 2018). These enhanced grafts may prove effective in some instances of neural regeneration, including promising results showing host regeneration following long gap repairs (Marquardt et al., 2015). However, despite these promising engineering feats, even with the addition of biologically-active filler, acellular grafts remain unable to address overall shortcomings in nerve regeneration, potentially due to limitations in the number of factors delivered, a non-optimal presentation of these factors, and/or lack of sufficient temporal and spatial control of their release. However, undesirable results have also been reported using over-engineered cellular grafts, providing a prime example of the “candy store effect” (Santosa et al., 2013). This effect was observed following the supplementation of NGTs with engineered Schwann cells over-expressing GDNF. Rather than promoting nerve regeneration with a strong chemoattractant for regenerating axons, the neurite outgrowth migrated to the site of GDNF release and did not move past this site of application – effectively countering the intended process of regeneration. As our data demonstrated, even by supplementing NGTs with disorganized DRGs, these grafts were still unable to provide signals sufficient to improve motor neuron cell health and support axon regeneration across the graft. Interestingly, others have shown that neurons directly transplanted into the distal nerve have improved muscle recovery in short-gap, long-gap, and delayed nerve repairs, possibly by providing direct structural integration and improving neurotrophic support (Kurimoto et al., 2016). However, just as advantageous structural attributes alone (e.g., alignment) are not enough for maximal regenerative capacity, a solely living component absent structural anisotropy also appears to be insufficient. For this reason, autografts remain the only readily accessible living scaffold strategy currently available in a surgeon’s armamentarium for peripheral nerve reconstruction that provides the native architecture and cellular support necessary for accelerating axonal regeneration, sustaining neuronal survival, and maximizing functional recovery. However, autografts have inherent shortcomings, including donor site morbidity and limited availability of donor nerve for long gap nerve repair and/or polytrauma resulting in multiple nerve injuries.

As a potential complimentary alternative to autografts, TENGs have continued to demonstrate the potential to overcome challenging limitations in nerve repair. Our group has previously reported that TENGs utilize a previously unknown mechanism described as “axon-facilitated axonal regeneration” to enable accelerated axonal regeneration and functional recovery compared to commercially-available NGTs, at levels equivalent to autograft repairs (Katiyar et al., 2020). Notably, TENG axons also penetrate the host distal nerve segment and “babysit” the distal Schwann cells as host regenerating axons cross the segmental defect – a mechanism not possible with autograft repairs. In the current study, we have shown TENGs also preserve the regenerative capacity of the proximal neuron populations within the spinal cord, thereby potentially increasing the ceiling for host regeneration and functional recovery (**Figure 1A**). In this study, TENG repairs resulted in a greater number of motor neurons maintained following PNI as compared to NGT repairs. In fact, within these motor neuron regions of the spinal cord, TENGs preserved FR and NeuN expression comparable to autograft repairs and naïve animals. Furthermore, we found a dramatic reduction in neuronal cell health following NGT repairs, which has been corroborated by clinical reports demonstrating poor functional recovery (Chang, Shah, Lee, & Yu, 2018). Although a qualitative assessment of functional recovery was performed to understand the chronic implications of the acute measures, further investigation is necessary to establish the importance of early preservation of proximal neurons and regenerative capacity on the rate and extent of functional recovery.

Although TENGs may represent a promising technology that preserves the regenerative capacity following repair and ultimately increases the ceiling for functional recovery, it will be necessary to test whether these findings persist in a large animal model with a cell source compatible for clinical translation. Indeed, the next phase of TENG development will involve the incorporation of a clinically-viable starting biomass, which may include neurons isolated from transgenic pigs lacking galactose-α1,3-galactose (gal) expression (Ekser et al., 2012). Traditional xenogenic tissue sources have historically been associated with risk for major postoperative complications resulting from tissue incompatibility due to ubiquitous gal expression. Previous studies have shown tissue derived from these transgenic pigs were compatible in primates. Therefore, “off the shelf” TENGs derived from this transgenic cell source may be suitable to act as a temporary living scaffold that facilitates axon regeneration and provides neurotrophic support for preserving the overall regenerative capacity of the patient.

## Conclusion

This study demonstrates that early surgical intervention with living scaffolds, such as autografts or TENGs, may preserve host spinal cord motor neuron health and regenerative capacity following acute segmental nerve repair. By preserving the regenerative capacity, living scaffolds may increase the potential ceiling for functional recovery compared to nonliving scaffolds. Therefore, TENGs may represent a promising technology for peripheral nerve repair by facilitating regeneration across segmental defects and providing crucial acute neurotrophic support necessary for successful functional recovery without the need for sacrificing an otherwise healthy donor nerve.

## Author Contributions

D.K.C. conceived the study and provided the experimental design. K.S.K. fabricated the TENGs. J.C.M. and J.C.B. performed the surgeries. K.D.B., Z.S.A., H.M.K., and J.M.R. provided critical perspective on the surgical paradigm, outcome measures, and interpretation. J.C.M., J.C.B., F.A.L., and K.D.B. conducted the histological assessments and analyses. J.C.M., J.C.B., and F.A.L. assisted with figure preparation. J.C.M., J.C.B., and D.K.C. prepared the manuscript. All authors provided critical feedback on the manuscript.

## Competing Financial Interests

D.K.C is a co-founder and K.S.K. is currently an employee of Axonova Medical, LLC, which is a University of Pennsylvania spin-out company focused on translation of advanced regenerative therapies to treat nervous system disorders. Multiple patents relate to the composition, methods, and use of tissue engineered nerve grafts, including U.S. Patent 9,895,399 (D.K.C.), and U.S. Provisional Patent 62/569,255 (D.K.C). No other author has declared a potential conflict of interest.

## Acknowledgements

The authors would like to thank Michael Johnson and Tom Villani of Visikol for technical support. Financial support provided by the U.S. Department of Defense [CDMRP/JPC8-CRMRP W81XWH-16-1-0796 (Cullen) & MRMC W81XWH-15-1-0466 (Cullen)], the Department of Veterans Affairs [BLR&D Merit Review I01-BX003748 (Cullen)], the National Institutes of Health [BRAIN Initiative U01-NS094340 (Cullen) & NRSA Graduate Research Fellowship F31-NS090746 (Katiyar)], and the Center for Undergraduate Research and Fellowships at the University of Pennsylvania. Opinions, interpretations, conclusions and recommendations are those of the author(s) and are not necessarily endorsed by the Department of Defense, the Department of Veterans Affairs, or the National Institutes of Health.

---

**Supplemental Figure 1.**
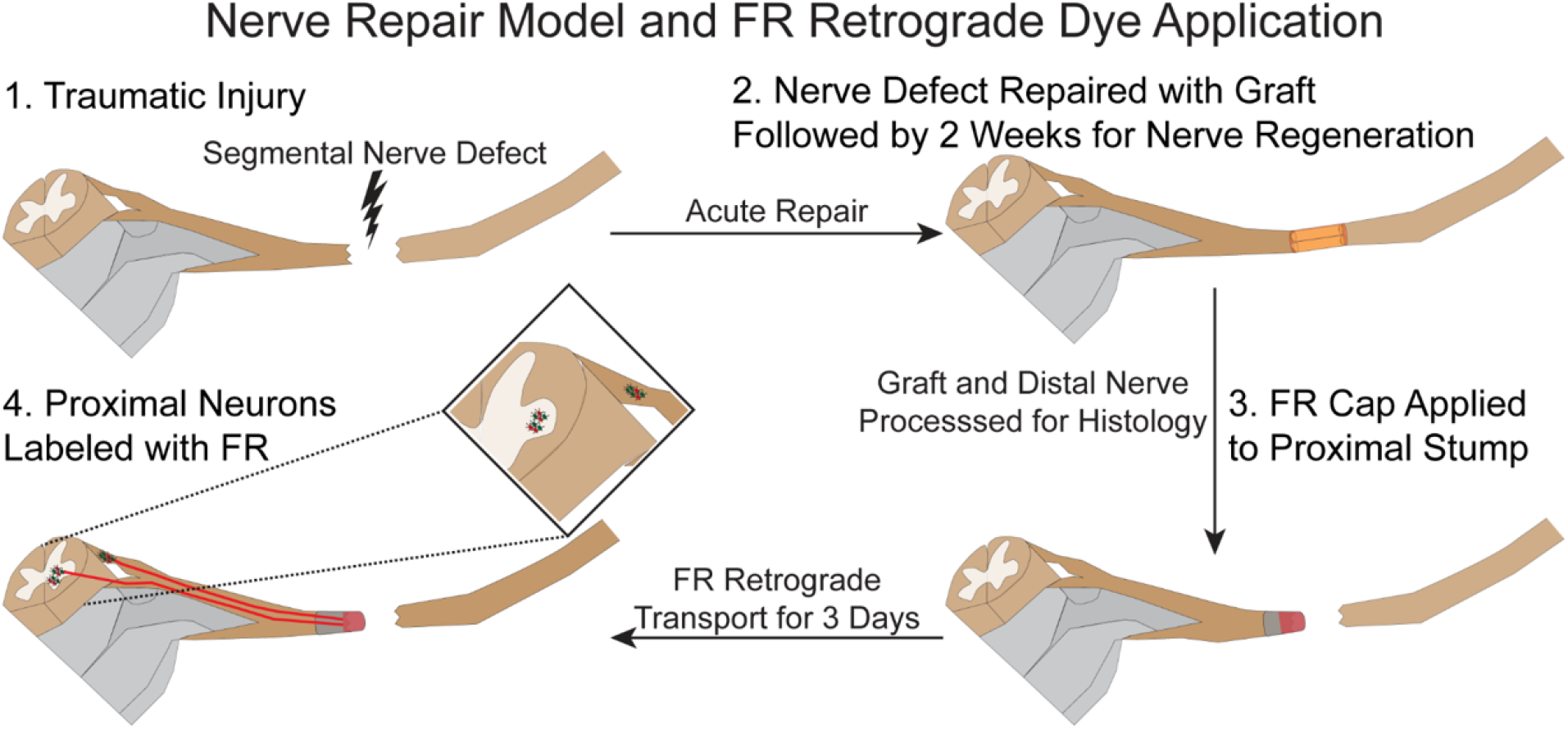
Fluoro-Ruby (FR) Application and Retrograde Transport. In this study, a retrograde fluorescent dye (FR) was applied at 2 weeks following repair of a 1 cm segmental nerve repair in a rat model. FR was applied proximal to the graft site, and the nerve graft zone (distal) was harvested for histological analysis. At 3 days post FR application, the animal was euthanized and the spinal cord and DRG were harvested for histological analyses.

**Supplemental Figure 2.**
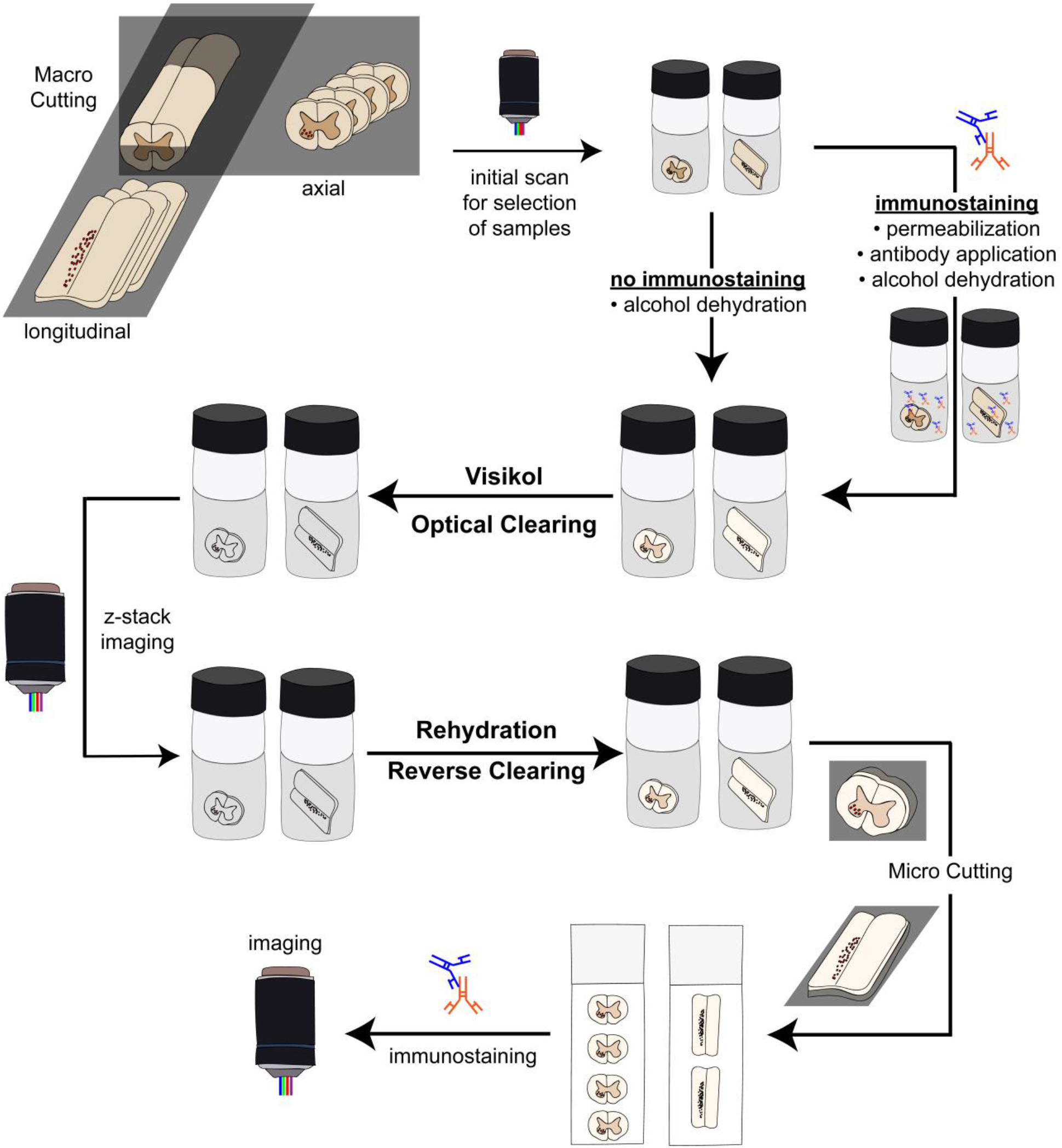
Spinal Cord Tissue Clearing, Staining, and Imaging Workflow. The surgical site was re-exposed at the terminal time and the graft was removed for further histological analysis. Fluororuby was applied to the proximal nerve stump and the animal was returned to the colony for an additional three days. Following transcardial perfusion, the spinal cord was harvested and blocked for optical clearing. After identifying the labeled tissue blocks, a subset were stained, optically cleared, and imaged using confocal microscopy. Positively labeled cells were then quantified from z-stack max projections. A subset of optically-cleared samples were reverse cleared for validation compared to traditional sectioning and quantification.

**Supplemental Figure 3.**
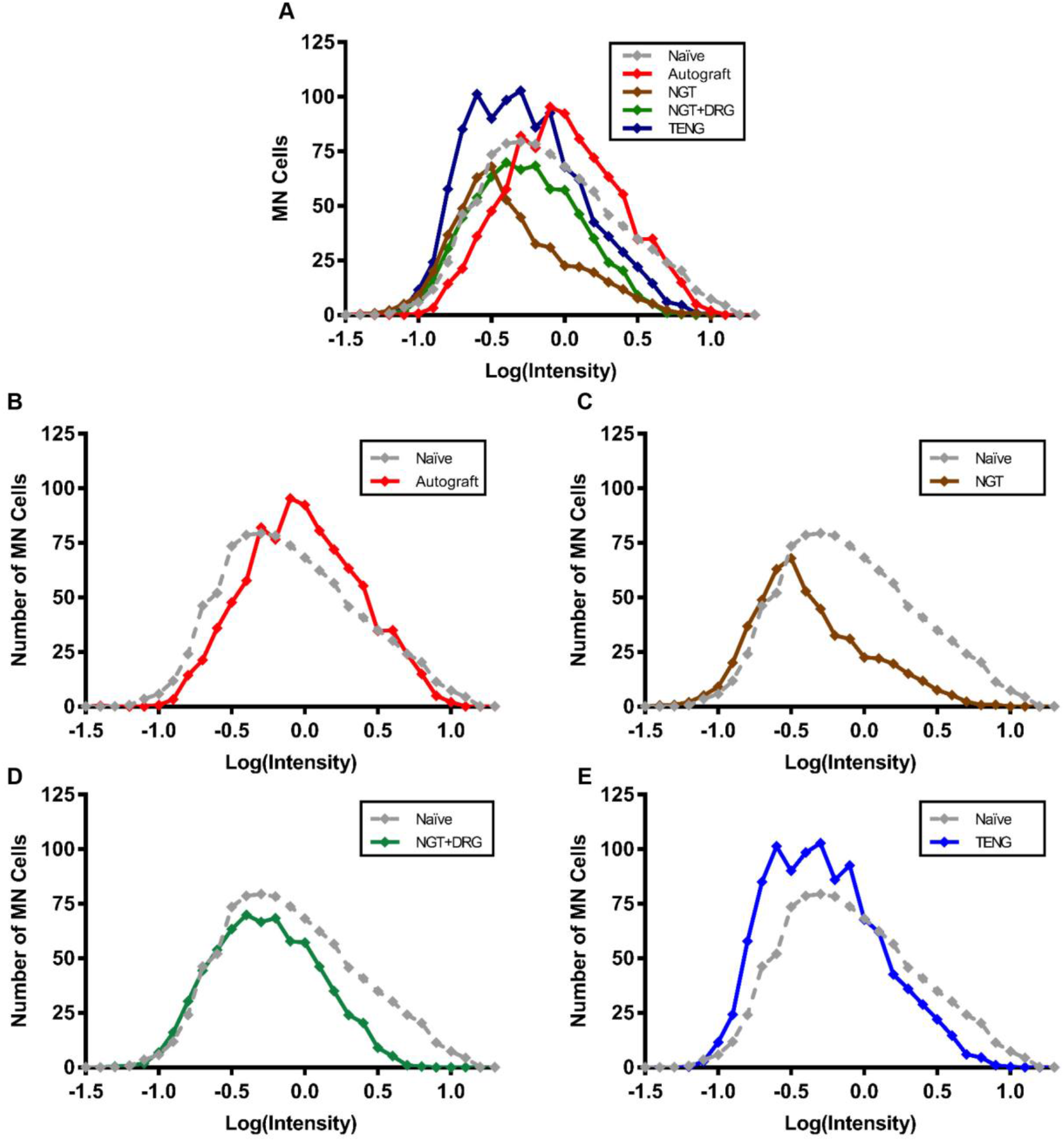
FR Intensity in Ventral Horn Motor Neuron Population. (A) Frequency distributions of MN fluorescence intensity were plotted. Intensity of FR fluorescence was calculated for each FR^+^ cell using maximum intensity of the cell relative to the local background around the cell. Individual cell intensity was log transformed to fit a normal distribution. (B-E) Frequency distributions for each experimental group were compared to the naïve frequency distribution. Note (A) is reproduced from **Figure 4** for convenience.

**Supplemental Figure 4.**
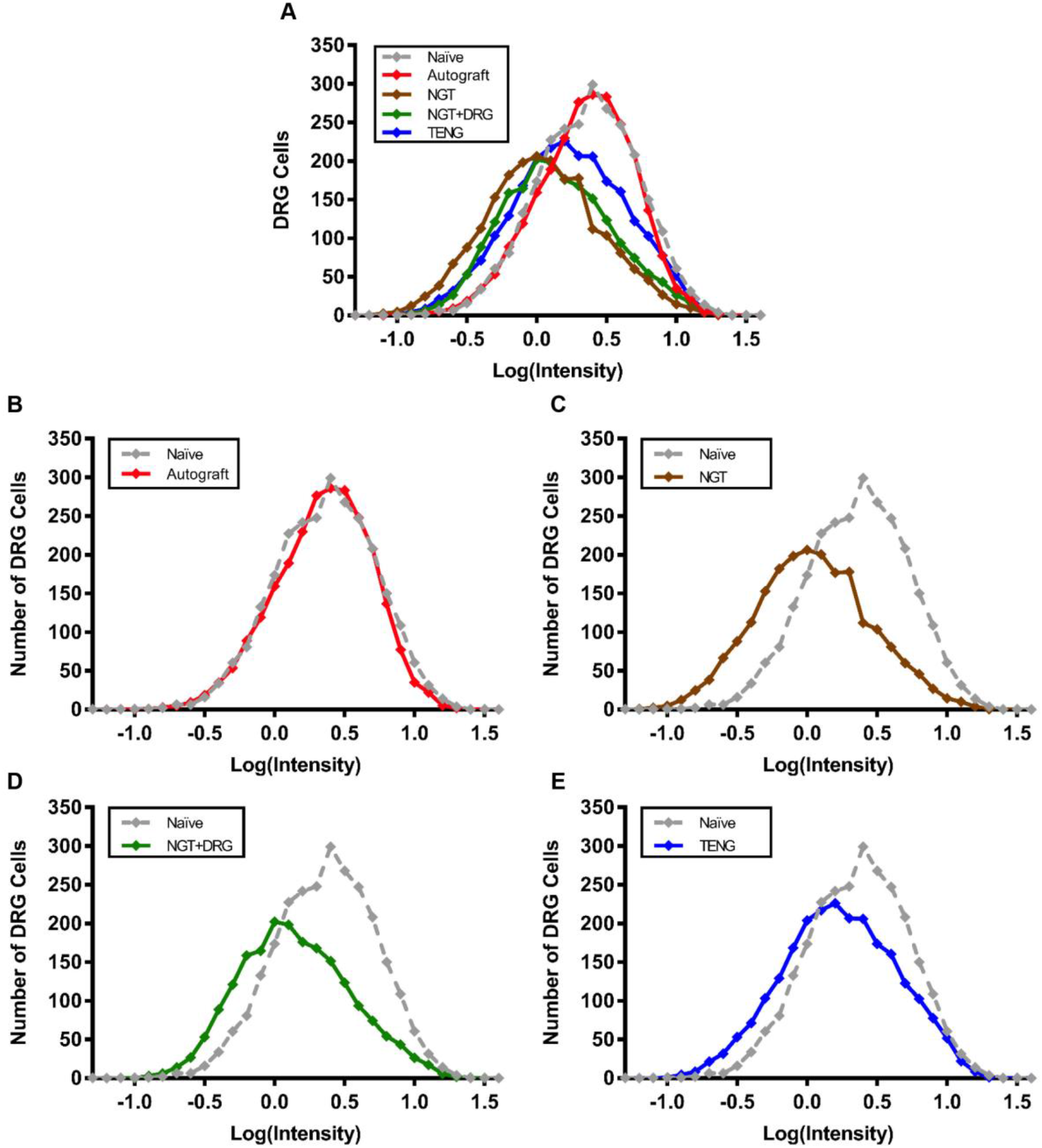
FR Intensity in L4 DRG. (A) Frequency distributions of DRG fluorescence intensity were plotted. Intensity of FR fluorescence was calculated for each FR^+^ cell using maximum intensity of the cell relative to the local background around the cell. Individual cell intensity was log transformed to fit a normal distribution. (B-E) Frequency distributions for each experimental group were compared to the naïve frequency distribution. Note (A) is reproduced from **Figure 5** for convenience.

